# Molecular events accompanying aggregation-induced energy quenching in Fucoxanthin-Chlorophyll Proteins

**DOI:** 10.1101/2024.03.20.585890

**Authors:** Maxime T. A. Alexandre, Tjaart P.J. Krüger, Andrew A. Pascal, Vasyl Veremeienko, Manuel Llansola-Portoles, Kathi Gundermann, Rienk van Grondelle, Claudia Büchel, Bruno Robert

**Affiliations:** Institute of Integrative Biology of the Cell, CEA, CNRS, Université Paris-Saclay, Avenue de la Terrasse, 91198 Gif sur Yvette, France; Department of Physics and Astronomy, Faculty of Sciences, VU University Amsterdam, De Boelelaan 1081 HV Amsterdam, The Netherlands; Department of Physics, Forestry and Agricultural Biotechnology Institute (FABI), Faculty of Natural and Agricultural Sciences, University of Pretoria, Private bag X20, Hatfield, 0028, South Africa; Institute of Molecular Biosciences, Department of Biosciences, Goethe University Frankfurt, Max von Laue Str. 9, 60438 Frankfurt, Germany

**Keywords:** Diatoms, Light-Harvesting, Xanthophyll, NPQ, Raman, Photoprotection

## Abstract

In high light, the antenna system in oxygenic photosynthetic organisms switches to a photoprotective mode, dissipating excess energy in a process called non-photochemical quenching (NPQ). Diatoms exhibit very efficient NPQ, accompanied by a xanthophyll cycle in which diadinoxanthin is de-epoxidized into diatoxanthin. Diatoms accumulate pigments from this cycle in high light, and exhibit faster and more pronounced NPQ. The mechanisms underlying NPQ in diatoms remain unclear, but it can be mimicked by aggregation of their isolated light-harvesting complexes, FCP (fucoxanthin chlorophyll-a/c protein). We assess this model system by resonance Raman measurements of two peripheral FCPs, trimeric FCPa and nonameric FCPb, isolated from high- and low-light-adapted cells (LL, HL). Quenching is associated with a reorganisation of these proteins, affecting the conformation of their bound carotenoids, and in a manner which is highly dependent on the protein considered. FCPa from LL diatoms exhibits significant changes in diadinoxanthin structure, together with a smaller conformational change of at least one fucoxanthin. For these LL-FCPa, quenching is associated with consecutive events, displaying distinct spectral signatures, and its amplitude correlates with the planarity of the diadinoxanthin structure. HL-FCPa aggregation is associated with a change in planarity of a 515-nm-absorbing fucoxanthin, and, to a lesser extent, of diadinoxanthin. Finally, in FCPb, a blue-absorbing fucoxanthin is primarily affected. FCPs thus possess a plastic structure, undergoing several conformational changes upon aggregation, dependent upon their precise composition and structure. NPQ in diatoms may therefore arise from a combination of structural changes, dependent on the environment the cells are adapted to.

## Introduction

During the photosynthetic process, light energy is absorbed by antenna proteins and efficiently transferred, in the form of an electronic excitation, to the reaction centers where photochemistry takes place. In low light conditions, the antenna system of plants and algae performs these energy transfer steps with very high efficiency. If the light intensity increases dramatically, such that the energy absorbed by the organism can no longer be fully transduced into chemical potential energy, the light-harvesting system switches into a photoprotective mode, dissipating the excess energy as heat^1^. This process is referred to as non-photochemical quenching of chlorophyll fluorescence (NPQ). In higher plants the major NPQ component (referred to as qE, for energy-dependent quenching) is triggered by the build-up of the pH gradient across the thylakoid membrane, and is facilitated by the de-epoxidation of violaxanthin to zeaxanthin in the xanthophyll (or more precisely, *violaxanthin*) cycle^2^. qE occurs in seconds, and so must correspond to a molecular re-organization of the existing photosynthetic apparatus. While it has been the object of discussion over many years, it is now generally agreed that the major light-harvesting protein LHCII of plants is the site of qE, and the formation of the quenching site involves a conformational change of this protein^3,4^. This conformational change drives the excitation energy in these proteins to the absorption-silent, lower-energy S_1_ excited state of one of the luteins bound to the complex^4^. An equivalent energy transfer process from chlorophyll (Chl) to the carotenoid (Car) S_1_ state was recently revealed in the permanently-quenched HLIP proteins in cyanobacteria^5^, which are considered the ancestors of the extended LHC family.

Diatoms are oxygenic photosynthetic organisms possessing an antenna system constituted of fucoxanthin chlorophyll-a/c proteins (FCP), members of the LHC family that differ from plant LHCs in both their pigment composition and organization. FCP binds a large number of the fucoxanthin (Fx) carotenoid, and contains Chl c instead of Chl b^6^. This pigment composition enhances the absorption cross-section in the blue-green region of the solar spectrum, and represents an adaptation of diatoms to their aquatic environment. The structures of a range of FCP complexes from different algae have recently been determined^7,8^. They are constituted of different oligomeric states of subunits with a structure closely related to that of the LHC monomer^7,8,9^.

Diatoms have to face enormous changes in light intensity when they are transported from deep water layers below the euphotic zone to full sunlight at the water surface. They possess very efficient photoprotective mechanisms to avoid photodamage, and NPQ appears to be more efficient in these organisms than in plants^10^. It comprises a fast component generated immediately upon the onset of light stress. This process is similar to qE in plants, depending on the ΔpH resulting from high-light exposure, and accompanied by a xanthophyll (*diadinoxanthin*) cycle in which diadinoxanthin (DD) is de-epoxidised to diatoxanthin (DT)^11,12,13^. While DT, like zeaxanthin in plants, may not be directly responsible for the quenching, its presence is nevertheless essential for the appearance of NPQ – an important distinguishing feature of the NPQ process in diatoms as compared to plants^14^. When diatoms are grown in high-light environments, the total amount of DD cycle pigments drastically increases, and the significantly higher amount of DT leads to a more pronounced NPQ which occurs more rapidly^10^.

The majority of the DD cycle pigments are located within FCP complexes^15,16^. In *Cyclotella meneghiniana*, two peripheral antenna complexes, the trimeric FCPa and nonameric FCPb, have been isolated^17,18^. FCPa and FCPb are mainly composed of the polypeptides Fcp6 & Fcp2 (now called Lhcx1 & Lhcf1/4), and Fcp5 (Lhcf3), respectively^17,19^. Analyses suggest the existence of a DD/DT-binding site in FCPa, while very little is found in FCPb preparations^20^. Carotenoids from the diadinoxanthin cycle were proposed to belong to three distinct pools, two of them bound to special antenna proteins within PSI and FCP, respectively, while an additional, much larger pool appearing in high-light-grown cells would be localized within a monogalactosyldiacylglycerol (MGDG) shield surrounding FCP (and co-purifying with it)^21^. However, selective observation of DD and DT *in vivo* by resonance Raman (RR) spectroscopy demonstrated that this additional pool of xanthophylls is also protein-bound^22^. As the de-epoxidation of DD is strongly enhanced in the presence of MGDG^23,24^, it is however possible that this non-bilayer lipid is involved in the binding of DD to protein. The multiple possible interactions of the complexes between themselves and with the lipids in the membrane create a vast binding site landscape for the extra carotenoids synthesized under HL conditions. Obviously, since these sites provide limited binding energy and are easily disrupted, they cannot be fully characterized in isolated complexes.

Molecular mechanisms involved in NPQ in diatoms are still not completely elucidated, and in particular the mode of action of DT. The presence of DT in isolated FCPa complexes appears to induce some limited quenching^20,25^, but the build-up of a pH gradient is mandatory to fully activate its quenching ability *in vivo*, possibly through the protonation of specific FCP residues^25,26^. Two independent quenching sites have been proposed to exist in *C. meneghiniana*^27^, one located in aggregated FCPs, and the second in PSII-associated FCPs. According to this hypothesis, the activation of the second site only would be dependent on the amount of DT. As observed for plant LHCs, the fluorescence of isolated FCPa aggregates exhibits a short lifetime and red-shifted emission^28,29^, and so they may constitute a model to study fluorescence quenching in diatoms. Previously, we characterized by resonance Raman the amount and conformation of the xanthophylls DD and DT *in vivo*, for whole cells in different light conditions^22^. Here we apply this method to FCPs, isolated from *C. meneghiniana* cells grown in different light environments and present in different aggregation states, to address the quenching mechanism in this *in vitro* model of NPQ.

## Experimental procedures

### Cell growth

The diatom *C. meneghiniana* (Culture Collection Göttingen, strain 1020-1a) was grown in batch culture at 18 °C, with constant shaking at 120 rpm. Cells were cultured according to Provasoli et al. 1957^30^, supplemented with 2 mM of silica, under low-light (40 μmol photons.m^−2^.s^−1^ (µE.m^−2^.s^−1^); LL) or high-light (140 µE.m^−2^.s^−1^; HL) regimes, having a 16 h light / 8 h dark cycle. These two light regimes have been shown to induce significant differences in various physiological parameters, including DD/DT content^31^.

### Purification of FCP

Cells were harvested in the early light phase by centrifugation after 7 days of culturing. Thylakoid membranes were isolated by several centrifugation steps after breaking the cells in a bead mill, according to Büchel (2003)^9^, except that the final buffer was complemented with EDTA (10 mM MES, 2 mM KCl, 5 mM EDTA pH 6.5, according to Beer et al., 2006^17^). Thylakoids were solubilised for 20 min on ice at 0.25 mg mL^−1^ total Chl a using 20 mM ß-dodecyl maltoside (βDM) (1:71 mol/mol). The complexes were then purified by ion exchange chromatography, followed by sucrose density centrifugation using a stepwise gradient in buffer 1 (25 mM Tris, 2 mM KCl, 0.03% βDM (w/v), pH 7.4) according to Gundermann and Büchel 2012^20^. The absorption spectra of the obtained complexes were similar to those in Ref.^17^. After preparation, FCP complexes were washed and concentrated using filtration devices with 30 kDa cutoff (Centripreps). To induce their aggregation, 250 μl of sample was either dialysed against 500 ml buffer without detergent (25 mM MES pH 5 or Tris pH 7.4) for 36 h, replacing the buffer after the first 12 h, or was incubated with Biobeads (Biorad). The quenching factor was determined by measuring their fluorescence emission spectra upon excitation at 440 nm.

### HPLC analysis

Pigment stoichiometries were determined by analytical HPLC (Merck Elite LaChrom, L-2130/L-2450, Germany). Cells were harvested by a short spin, freeze-dried, re-suspended in 90 % acetone, 10 mM Tris pH 7.5, homogenised in a tissuelyser (Quiagen), and sonicated for 10 min (Elmasonic, Elma). Samples were then centrifuged shortly at maximum speed in an Eppendorf centrifuge, and the supernatants were loaded onto the column. Pigment extraction from isolated complexes was performed using the same acetone concentration followed by centrifugation. After separation on a RP18 monolith column, pigments were quantified using a photodiode array detector. Calibration and quantification were carried out as described in ^32^.

### Resonance Raman spectroscopy

A drop of purified FCP was deposited on a microscope coverslip, immediately immersed in liquid nitrogen (LN_2_) and transferred into an LN_2_-flow cryostat (Air Liquide). Raman scattered photons were collected at 90° to the incident beam and focused into a U1000 double-monochromator Raman spectrometer (Jobin-Yvon) equipped with a back-illuminated, LN_2_-cooled charge-coupled-device camera. The resulting spectral resolution is below 2 cm^−1^, with a wavenumber accuracy, after alignment on the Rayleigh band (at 0 cm^−1^), of better than 1 cm^−1^. Laser excitations were obtained using an Ar^+^ laser (Sabre, Coherent).

## Results

### Pigment composition of isolated FCP

Since Beer et al. 2006^17^, the isolation of FCPa and FCPb has been significantly improved. Premvardhan et al. 2009^33^ concluded from HPLC analysis that FCPa binds more DD than FCPb, in line with the previous work of Beer et al.^17^. Table 1 summarises the pigment content of the preparations used here, together with that of the cells they were extracted from. Improvements in the isolation procedures lead nowadays to FCP complexes binding significantly more Fx. FCPa extracted from HL cells binds twice as much DT as the same complex obtained from LL cells, and four times more DD than FCPb extracted from the same cells (the carotenoid composition of the latter being much less sensitive to the growth light environment).

**Table 1:** HPLC pigment content (± standard deviation; n= 3) of *Cyclotella* cells grown in different light conditions, and of the purified FCPs from these cells, expressed in mole per mole of Chl-a.

| | Chl c | Fx | DD | DT | β-Car |
| --- | --- | --- | --- | --- | --- |
| HL-Cells | 0.153 ($\pm 0.001$) | 0.598 ($\pm 0.006$) | 0.316 ($\pm 0.010$) | 0.088 ($\pm 0.003$) | 0.052 ($\pm 0.000$) |
| LL-Cells | 0.160 ($\pm 0.002$) | 0.589 ($\pm 0.01$) | 0.119 ($\pm 0.000$) | 0.020 ($\pm 0.000$) | 0.044 ($\pm 0.002$) |
| HL-FCPa | 0.311 ($\pm 0.000$) | 0.988 ($\pm 0.005$) | 0.024 ($\pm 0.001$) | 0.018 ($\pm 0.000$) | 0.006 ($\pm 0.001$) |
| LL-FCPa | 0.318 ($\pm 0.007$) | 1.002 ($\pm 0.003$) | 0.021 ($\pm 0.000$) | 0.009 ($\pm 0.000$) | 0.005 ($\pm 0.000$) |
| HL-FCPb | 0.333 ($\pm 0.003$) | 0.897 ($\pm 0.007$) | 0.006 ($\pm 0.001$) | 0.007 ($\pm 0.000$) | 0.002 ($\pm 0.000$) |
| LL-FCPb | 0.326 ($\pm 0.002$) | 0.890 ($\pm 0.002$) | 0.008 ($\pm 0.001$) | 0.005 ($\pm 0.001$) | 0.002 ($\pm 0.000$) |

### In vitro model for quenching: aggregation of FCP complexes

We induced aggregation of the FCP complexes by detergent removal, either by dialysis or by using Biobeads. Aggregation induces a dramatic decrease in the fluorescence of these proteins – reported in Table 2 are the quenching factors achieved, corresponding to the fluorescence intensity ratio between the disaggregated and aggregated protein. The extent of quenching of FCPa induced by aggregation did not depend on the light environment of the cells the complexes were initially extracted from, and reached 7-8 or 12-13 upon dialysis at pH 7.4 or 5.0, respectively, and 20 when removing detergent with Biobeads – values which are consistent with previous reports^27^. FCPb aggregation generally induces much more limited quenching, of 4 upon dialysis (irrespective of pH) and 6 upon Biobead treatment (Table 2).

**Table 2:** Quenching factors observed for the different FCP preparations, according to the aggregation procedure, calculated as (F-disaggregated / F-aggregated) where F is the integrated area of the fluorescence peak in each case.

|  | Dialysis pH 7.4 | Dialysis pH 5 | Biobeads |
| --- | --- | --- | --- |
| HL-FCPa | 8 | 13 | 20 |
| LL-FCPa | 7 | 12 | 20 |
| FCPb (HL or LL) | 4 | 4 | 6 |

### Resonance Raman spectra of the carotenoid molecules bound to FCP

Carotenoid resonance Raman (RR) spectra generally contain four groups of bands, termed ν_1_ to ν_4_. The ν_1_ and ν_2_ bands arise from stretching modes of the conjugated C=C and C-C bonds, and are located around 1525 and 1160 cm^−1^, respectively^34^. Their position and shape are slightly different for Fx than for DD or DT^22^. The ν_3_ band, which contributes at 1000-1020 cm^−1^, arises from in-plane rocking vibrations of the methyl groups attached to the conjugated chain, coupled with in-plane bending modes of the adjacent C-H’s^34^. It was recently shown that it can be used as a fingerprint of the configuration of conjugated end-cycles^35^, and the presence of allene and alkyne groups in Fx and DD/DT, respectively (see Fig. S1a), is probably responsible for their significant variation from more “classical” linear carotenoids, as well as from each other^36,37^. The ν_4_ band located around 960 cm^−1^ arises from C-H out-of-plane wagging motions coupled with C=C torsional modes (out-of-plane twists of the carbon backbone)^34^. These out-of-plane modes are not resonance-enhanced when the molecule is planar, but gain amplitude upon out-of-plane distortions of the carotenoid structure. It thus reflects the degree of twisting of these molecules. Resonance Raman spectra of Fx, DD and DT in the ν_3_ and ν_4_ regions are presented in Figure 1.

**Figure 1:**
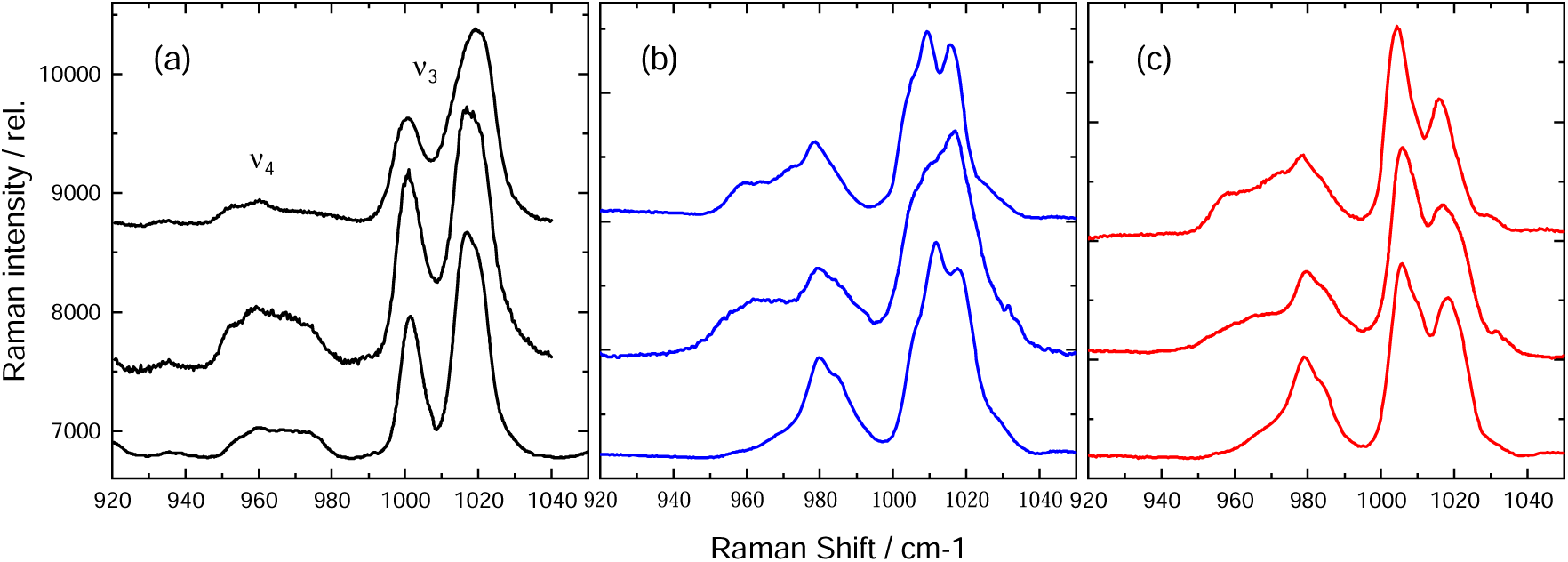
ν_3_ and ν_4_ regions of resonance Raman spectra obtained at 77 K of: fucoxanthin (a) in diethyl-ether, dichloromethane and pyridine (from top to bottom), and of diadinoxanthin (b) & diatoxanthin (c) in toluene, pyridine and ethanol. Excitations to induce resonance were chosen close to the absorption maximum in each case, at 488.0 nm for fucoxanthin, 496.5 nm for diadinoxanthin and 514.5 nm for diatoxanthin (see absorption spectra in Fig. S1b).

In the ν_3_ region, RR spectra of Fx (Fig. 1a) display two well-separated bands at 1000-1002 and 1018-1020 cm^−1^. Spectra of DD (Fig. 1b) contain convoluted components, forming a broad cluster with a maximum peak at *ca* 1010-1012 cm^−1^. The spectra of DT (Fig. 1c) generally exhibit an intense band at 1004-1007 cm^−1^, accompanied by a ∼30 % weaker component at 1016 cm^−1^. The relative intensity of these bands varies depending on the solvent, but this region can still be used as a fingerprint to determine which of these carotenoids contribute to a given Raman spectrum. In the ν_4_ region, spectra of all three isolated carotenoids contain modes indicating that their equilibrium structure is somewhat distorted, and that this distortion is solvent-dependent. Of course, the modes in this region will be sensitive to, and actually determined by, the steric constraints imposed on the carotenoids by their protein binding pocket.

In the following, we have analysed the resonance Raman spectra of FCPa and FCPb complexes, and how these spectra evolve when the proteins aggregate. Resonance Raman spectra were measured using excitation wavelengths throughout the absorption band of the FCP-bound carotenoids, namely at 488.0, 496.5, 501.7 and 514.5 nm. At all these excitation wavelengths, all carotenoids present in the complexes should contribute to the Raman spectra, although with different relative intensities.

### Effect of LL-FCPa aggregation: resonance Raman spectra at 496.5 nm excitation

The most significant Raman changes upon aggregation of LL-FCPa were observed using 496.5 nm excitation. Figure 2 displays the FCPa spectra in these excitation conditions, for a range of aggregation states. LL-FCPa aggregation induces an alteration mainly affecting the ν_3_ and ν_4_ regions. As the carotenoid content of FCPa is unchanged during aggregation (as observed by HPLC analysis; data not shown), these spectral changes must arise either from changes in resonance conditions (*i.e.* a change in the relative contribution of different carotenoid populations) or from structural alterations to the contributing carotenoids. Changes in resonance should affect the whole spectrum, but no significant evolution of the ν_1_ and ν_2_ bands is observed (Fig. 2b). The absence of changes in these latter regions thus allows aggregation-induced changes in resonance to be ruled out.

**Figure 2:**
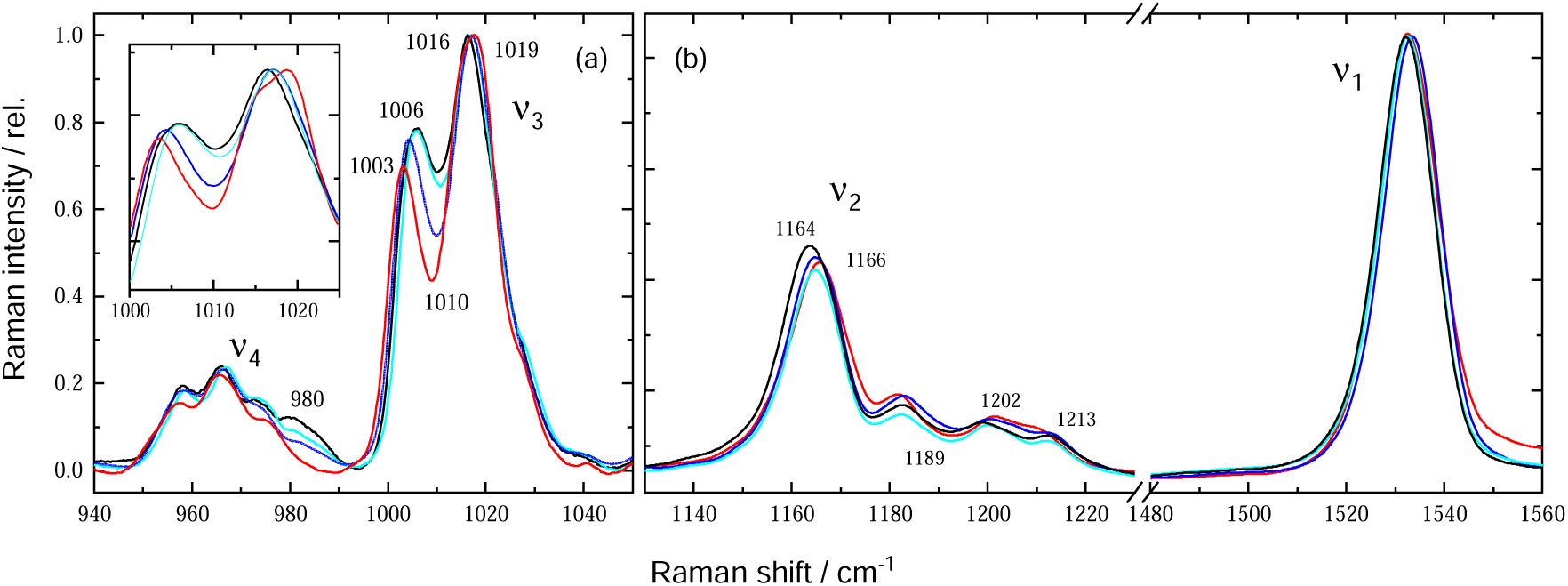
Resonance Raman spectra excited at 496.5 nm of trimeric LL-FCPa (black), and upon aggregation by dialysis at pH 7.4 (cyan) & pH 5.5 (blue), or using Biobeads (red). Spectra in panel a (ν_3_ & ν_4_) have been multiplied by 6.5 relative to panel b (ν_1_ & ν_2_) for clarity; normalisation was performed separately for panels a & b (using ν_3_ & ν_1_ peaks, respectively). Inset: magnification of the ν_3_ bands (1000-1030 cm^−1^).

A clear decrease in the intensity of the ν_4_ component at 980 cm^−1^ is observed upon FCPa aggregation, evident already when NPQ reaches 7 (dialysis at pH 7.4; cyan spectrum). At higher NPQ, a concomitant distortion of ν_3_ is observed, together with a decrease in the 960, 975 and 980 cm^−1^ components of ν_4_. This suggests the presence of at least two different mechanisms upon successive FCPa aggregation. Firstly, at NPQ=7, a carotenoid population has become more planar. At higher NPQ upon further aggregation, either this population gets even more planar or a second population is affected. The spectral changes in the ν_3_ region are complex. FCPa aggregation obtained after dialysis against buffer at pH 7.4 induces a slight upshift in the higher-frequency component, from 1016.0 to 1016.5 cm^−1^, while the lower-frequency component remains unchanged. Higher NPQ values (dialysis at pH 5.5 or Biobead treatment) induce a further upshift in the higher-frequency component (up to 1019 cm^−1^ with Biobeads), a significant downshift of the lower-frequency component (from 1006 to 1003 cm^−1^), and a net decrease in the intensity of the signal at 1010 cm^−1^. A more thorough analysis of the evolution of these spectra is shown in the Supplementary Material (Figs S2 & S3 and accompanying text), and reveals that at least four consecutive spectral events (and thus five distinct LL-FCPa configurations) are necessary to explain the evolution of these spectra.

It is worth noting that a clear correlation is observed between the quenching values obtained upon LL-FCPa aggregation and the intensity changes in the 980 cm^−1^ component of ν_4_ (Fig. 3).

**Figure 3:**
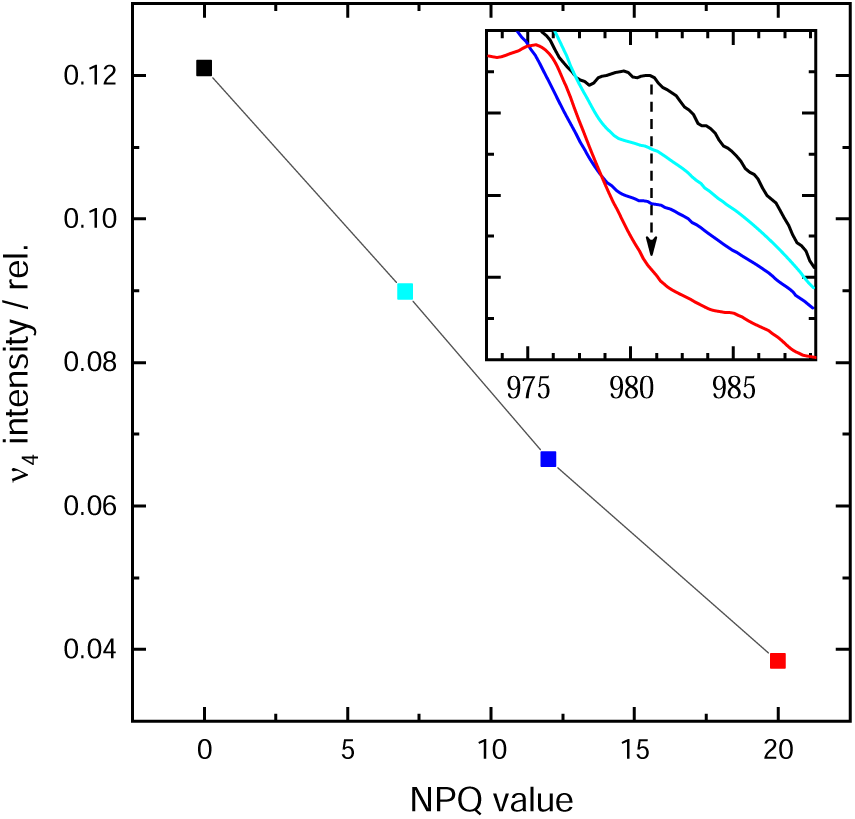
Correlation between the intensity at 981 cm^−1^ and the NPQ value for successive LL-FCPa aggregation. Inset: high-frequency component of the ν_4_ region for LL-FCPa trimers, and upon successive aggregation (colours as per Fig. 2).

### Identification of the carotenoids altered during aggregation of LL-FCPa

In whole cells, the concentration of Fx is two to six times higher than that of DD and DT (Table 1), but excitations between 476 and 514 nm *in vivo* yield RR spectra dominated by contributions of the latter (DD cycle) carotenoids^22^. This was attributed to the fact that the absorption spectra of the Fx molecules are spread over a very large spectral range, while those of DD and DT molecules are far less inhomogeneously broadened, such that the DD cycle carotenoids exhibit a larger resonance cross-section in this region^22^. In isolated FCPa complexes extracted from LL cells, the relative amount of DD is fivefold less than for whole cells (Table 1), and a much lower contribution of DD is expected. However, contributions from Fx only cannot fully account for the spectra obtained at 496.5 nm. The shape of ν_3_ (compare with Figure 1) clearly reveals the presence of additional contributions, which, in the absence of DT, must arise from DD. This is confirmed for this excitation when comparing spectra of LL- and HL-FCPa, which exhibit about the same Fx/DD ratio, to those of FCPb, binding three times less DD (Figure 4). At this excitation, the ν_3_ region for FCPb exhibits two well-separated components, at *ca* 1000 and 1020 cm^−1^, with a gap at 1010 cm^−1^, similar to those observed for isolated Fx (Fig. 1a). In contrast, these bands are highly convoluted in FCPa spectra, observable as a smaller 1010 cm^−1^ trough between the two main peaks, confirming the mixed contributions of Fx and DD. The fact that the component sensitive to aggregation lies in between the two main bands (see Fig. 2) strongly suggests that the changes observed mainly arise from DD bound to these complexes.

**Figure 4:**
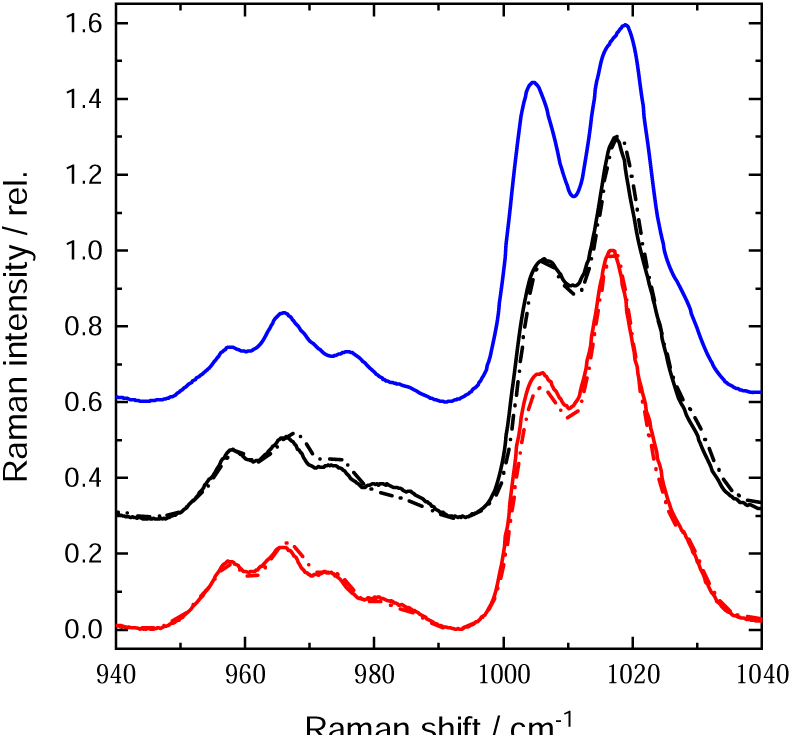
ν_3_ and ν_4_ regions of RR spectra at 496.5 nm, shown for FCPb (blue), LL-FCPa (black), and HL-FCPa (red). Aggregation (dotted lines) was induced by dialysis at pH 7.4 (not shown for FCPb).

Although the calculated cross-correlation between the four resonance Raman spectra in Fig. 2a reveals that at least five different spectral states are necessary to account for all the variations observed during aggregation of LL-FCPa (Fig. S3), we cannot extract the spectral signatures of each of these steps from this set of data. In the absence of such information, we can only conclude that at least four independent structural events occur upon FCPa aggregation, two of them arising from changes in configuration affecting DD molecules, and the other two possibly arising from DD or Fx molecules.

### Effect of HL-FCPa aggregation: resonance Raman spectra at 496.5 nm

A similar analysis was performed for aggregation of HL-FCPa. Figure 4 compares the effect of aggregation of LL- and HL-FCPa, induced by dialysis at pH 7.4, on their RR spectra at 496.5 nm excitation. The decrease in intensity of the ∼980 cm^−1^ band is smaller in HL-FCPa compared to the changes seen in LL-FCPa, and accompanied by even smaller changes in the other components of the ν_4_ region.

FCPa extracted from HL cells contains about 15 % more DD than LL-FCPa (Table 1). The poor sensitivity of the DD bound to these complexes towards aggregation suggests that these carotenoids are bound to a different binding pocket in HL-FCPa than in LL-FCPa, as proposed elsewhere^22^. The weak overall response of these spectra to aggregation further suggests that a large part of the DD found in LL-FCPa is either absent in HL-FCPa or, more probably, has been deepoxidised into DT, which is expected to contribute poorly to the spectra obtained at this excitation. The smaller decrease of the 980 cm^−1^ band observed for HL-FCPa indicates that a smaller population of DD undergoes a similar change as that described for LL-FCPa above. This subpopulation may either represent a small fraction of DD bound to HL-FCPa, or arise from a small population of complexes similar to LL-FCPa present in these growth conditions. Aggregation of HL-FCPa also induces a decrease in the intensity of the ν_3_ band, which is further explored below.

### Effect of FCPa aggregation at other excitation wavelengths

When the excitation is blue-shifted to 488.0 nm, LL-FCPa exhibits similar, though less intense, aggregation-induced changes in resonance Raman spectra to those observed at 496.5 nm, and only a weak decrease in the intensity of the ν_3_ band is observed for HL-FCPa (data not shown).

Given the differences in absorption between DD and DT, excitations above 500 nm are expected to enhance DT contributions (as well as Fx) – as observed in whole cells^22^. Shown in Figure 5a are RR spectra of LL- and HL-FCPa complexes for the red-shifted excitation of 501.7 nm. While the spectra are quite similar, the difference “HL-FCPa *minus* LL-FCPa” does indeed correspond to DT (blue trace in Fig. 5a; cf Fig. 1), reflecting a doubling in DT stoichiometry (Table 1). The aggregation-induced changes observed for LL-FCPa are very similar at this wavelength to those observed at 496.5 nm, though less intense. On the other hand, far more significant aggregation-induced changes are observed at 501.7 nm for HL-FCPa, with in particular a large reduction in the 1006 cm^−1^ band (see difference spectrum in Fig. S2, red dashed trace). This must reflect significant structural changes in the DT present in these complexes, probably around or near the end-rings (see above).

**Figure 5:**
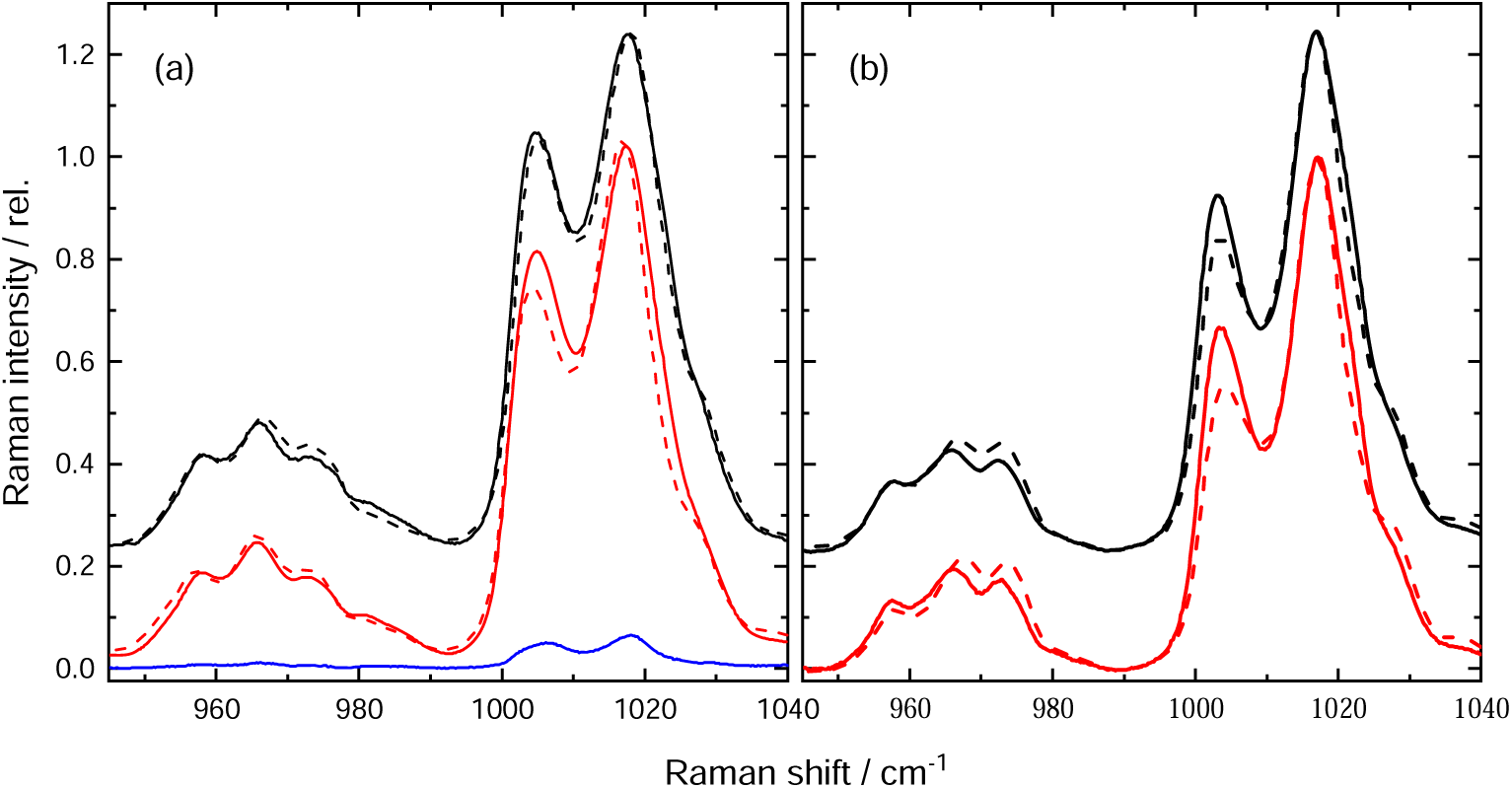
ν_3_ and ν_4_ regions of RR spectra, excited at 501.7 & 514.5 nm (a, b respectively), of LL-FCPa (black) and HL-FCPa (red) in the trimeric and aggregated states (solid, dotted lines, respectively). Difference “HL-FCPa *minus* LL-FCPa” is also shown at 501.7 nm (blue). Aggregation was induced by dialysis at pH 7.4.

Moving further to the red, Raman spectra obtained at 514.5 nm are almost identical for FCPa trimers extracted from cells exposed to different light environments (Fig. 5b), even though much less DT is bound to LL-FCPa. The contribution of DT to these spectra must therefore be weak. Moreover, the bands at 1000 and 1016 cm^−1^ are well-separated, indicating that DD contributions are also weak at best. Aggregation of these complexes induces a net decrease in intensity of the lower-frequency component of the ν_3_ band, accompanied by a tilt in relative intensities in the ν_4_ band, with the higher-frequency component increasing while the lower-frequency one decreases (difference spectrum reported in Fig. S2, black dashed trace).

The observed changes at 514.5 nm must originate from a population of Fx (the only carotenoid contributing to these spectra; see above). In HL-FCPa, this Fx population undergoes larger changes, or a larger Fx population undergoes these changes, as compared to LL-FCPa. It must be noted that such large changes in intensity may (at least in part) account for the small changes in the ν_3_ region observed at 496.5 nm excitation in HL-FCPa, and thus may explain the differences discussed in the previous paragraph. In a similar way, this Fx configurational change could be at the origin of the third spectral event observed in LL-FCPa at 496.5 nm. If this is the case, it would indicate that Fx would change configuration in LL-FCPa at a different state of aggregation of the protein than those involving DD.

### Effect of FCPb aggregation

FCPb aggregation induces less pronounced fluorescence quenching, and the observed changes in RR spectra are concomitantly smaller. Only minor aggregation-induced changes can be observed for 496.5 or 514.5 nm excitation (not shown), but significant spectral alterations are seen for excitation at 488 nm (Fig. 6). At this excitation wavelength, a net decrease of the lower-frequency component of ν_3_ is observed, possibly with very small changes in the relative intensity of the ν_4_ components. These changes must be attributed to Fx, the only carotenoid present in significant amounts in this complex. They are somewhat reminiscent of those observed for FCPa at 514.5 nm excitation (Fig. 5). However, as they are observed at a different excitation wavelength (they are *less* well observed at 514.5 nm), they must arise from an Fx with an absorption peak located further to the blue.

**Figure 6:**
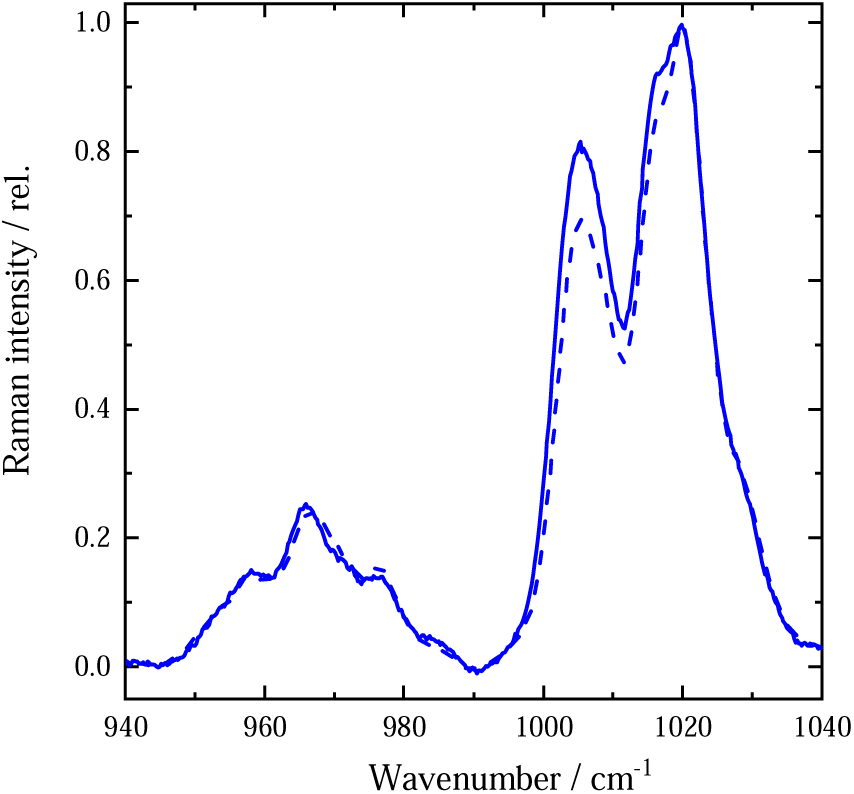
ν_3_ and ν_4_ regions of RR spectra of FCPb, obtained using 488.0 nm excitation, before and after aggregation by dialysis at pH 7.4 (solid, dotted lines, respectively).

## Discussion

From our RR results, it is possible to formally conclude that, as for LHCII, FCP complexes possess a plastic structure, undergoing several structural changes upon aggregation. Single molecule results have already been used to propose that FCPa possesses a more flexible structure than LHCII^38^. RR spectroscopy reveals these structural changes by their effect on the configuration of the protein-bound pigments. It should be stressed that these observations do not indicate which pigment(s) are directly responsible for quenching. (By comparison, in LHCII, the larger Raman change observed upon aggregation arises from the bound neoxanthin, but this carotenoid is not directly involved in the energy quenching process^3^.) Neither do they indicate that the structural changes in FCP are limited to these pigments. We thus obtain a partial view of what occurs to FCP upon aggregation, which nevertheless leads to a series of conclusions.

Aggregation of FCPa affects the configuration of at least three distinct carotenoid pools: one population each of diadinoxanthin, fucoxanthin and diatoxanthin. In FCPa extracted from cells grown in low light, the largest Raman change observed upon aggregation arises from the diadinoxanthin configuration, while Fx and DT are primarily affected in FCPa extracted from high-light cells. This suggests that the FCPa structure does not always respond in the same way to aggregation, and that this depends on the growth conditions of the cells. Such a conclusion is not as surprising as it may look, as the polypeptide composition of LL- and HL-FCPa is different, the latter comprising a larger proportion of Lhcx1^19^.

For LL-FCPa, a many-step process is observed upon progressive aggregation. Mild aggregation, leading to limited NPQ (quenching factor 7), induces a clear intensity decrease of the ν_4_ band, indicating that a diadinoxanthin population reaches a more planar structure. Upon further aggregation, the changes in ν_3_ indicate that the end-ring configuration of these carotenoids is modified, together with a further increase in planarity of these molecules. It is of note that a direct correlation is observed between the planarity of the diadinoxanthin and the extent of observed quenching along the whole aggregation process. 2D-correlation analysis of the RR spectra during this process further indicates that two additional carotenoid changes are observed, possibly affecting a pool of Fx.

The changes observed for DD in HL-FCPa are much smaller, in line with the conversion of DD to DT under this condition. However, some of these FCPa complexes may be similar to those present in LL cells. HL- and LL-FCPa show a different polypeptide composition, with HL-FCPa being enriched in Lhcx1 polypeptides, but only to a level that is most probably less than a third of the total^19^. This implies a mixture of different trimers in HL-FCPa, with a fraction of the complexes resembling LL-FCPa in their polypeptide composition. The limited diadinoxanthin changes observed could arise from this “LL-FCPa-like” subpopulation. On the other hand, a significant increase in the intensity of some components of the ν_4_ band is observed in HL-FCPa upon aggregation, which is not observed in LL-FCPa. As already stated, the obtained spectra do not arise from a single population of carotenoids. In HL-FCPa, an Fx configurational change is observed, inducing an increase in intensity of the same ν_4_ components. Although this is best observed for excitation at 514.5 nm, some remnants of this signal could well survive at 496.5 nm excitation. Therefore, we cannot formally conclude that the first step of HL-FCPa aggregation affects the configuration of a diadinoxanthin molecule. For these complexes, an additional out-of-plane torsion of a fucoxanthin population is clearly observed (as evidenced by the ν_4_ changes at 514.5 nm). The accompanying changes in the intensity of the ν_3_ components could again reflect a change in the configuration of the conjugated end-cycle of this carotenoid. This change is much weaker in LL-FCPa, possibly reflecting either the presence of a sub-population in these complexes, or a similar, but less extended structural change occurring upon their aggregation. On the other hand, the DT present in HL-FCPa also displays significant conformational changes upon aggregation, as seen at 501.7 nm. DT in LL-FCPa may also exhibit these changes, but its observation is at the limit of the measurement for the complex from low-light-grown cells.

In strong contrast to FCPa, aggregating FCPb leads to limited energy quenching only, even when using Biobeads. RR spectra of FCPb display limited changes upon mild aggregation, which are not followed by any additional observable changes upon further aggregation (data not shown). It is of note that these Raman changes are only observed upon mild aggregation, *i.e.,* during the step influencing their quenching state. In these conditions, a unique carotenoid structural change can be deduced from the RR spectra, probably concerning fucoxanthin absorbing around 490 nm, and showing that the configuration of this carotenoid pool is affected both in its planarity and at the level of its conjugated end-cycle.

If FCPa, like LHCII, displays a plastic structure which changes upon aggregation, and the changes correlate with their quenching ability, there is a major difference between these two classes of complex. In LHCII, no carotenoid from the xanthophyll cycle exhibits changes to its structure during aggregation. In FCPa extracted from low-light-grown cells, it is clear that the structural changes affect primarily diadinoxanthin, while for the same complex in high-light-grown cells the diatoxanthin is significantly affected. This may be considered consistent with the observation that diatoxanthin is an obligatory component of NPQ *in vivo*, unlike its counterpart zeaxanthin in plants. In FCPa extracted from high-light-grown cells, the diadinoxanthin structural changes are much weaker or even absent. However, it may be that the aggregation-sensitive DD has been converted into DT in these complexes.

The next steps in determining the state of FCP proteins in diatoms, and assessing any changes to this state associated with NPQ, will involve studies in more intact systems (isolated membranes, up to whole diatoms). These studies will be directed towards determining the structure and association state of these complexes in the intact membrane, and in confirming whether similar changes in pigment conformation are observed upon NPQ induction as seen here upon FCP aggregation *in vitro*. In higher plants, significant insight into the association of LHCs between themselves and with other complexes in the membrane have been achieved, through a combination of electron microscopy of isolated membrane fragments^39,40^, and sucrose gradient centrifugation to isolate macro-complexes of varying sizes^41,42^. At the same time, Raman measurements on membranes and leaves confirmed that a conformational change, similar to that observed upon LHCII aggregation *in vitro*, was intimately associated with NPQ in higher plants^4^. In the case of diatoms, on the one hand only a limited number of studies have been performed to determine the association state of FCPs *in vivo*^7,43^, but this type of approach is ongoing in a number of laboratories. On the second point, we will soon be in a position to publish data confirming that the conformational changes reported here *in vitro* are indeed observed in intact diatoms upon NPQ induction.

## Acknowledgments

Spectroscopic measurements were performed on the Biophysics Platform of I2BC, supported by the French Infrastructure for Integrated Structural Biology (FRISBI) ANR-10-INBS-05. CB acknowledges funding by the Deutsche Forschungsgemeinschaft (DFG Bu812/10)

