## Supplementary Material for "Molecular events accompanying aggregation-induced energy quenching in Fucoxanthin-Chlorophyll Proteins"

### Chemical structures and absorption spectra of FCP carotenoids

Chemical structures of the FCP carotenoids fucoxanthin, diadinoxanthin and diatoxanthin are given in Figure S1a, along with their absorption spectra in Fig. S1b.

(a)

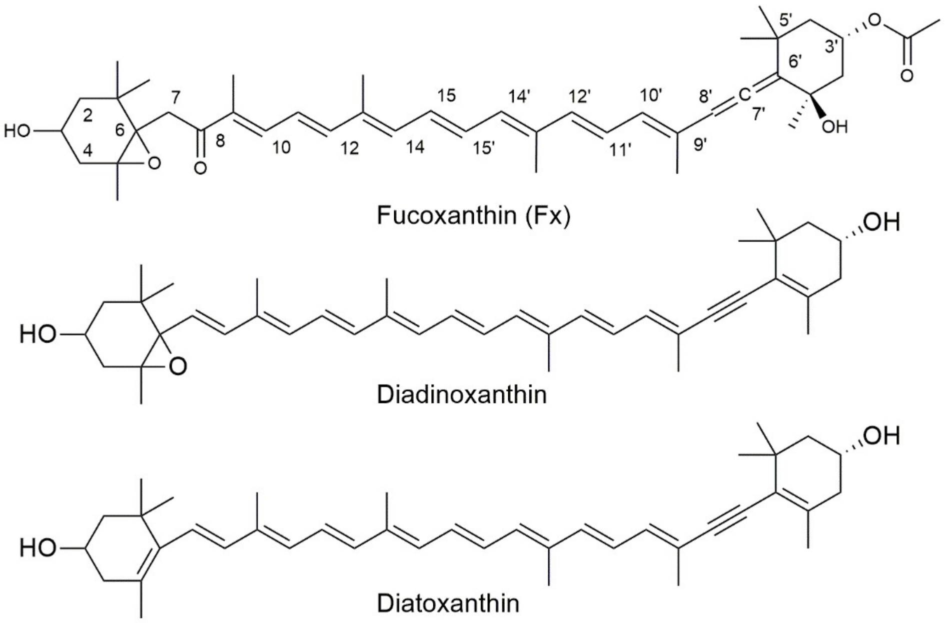

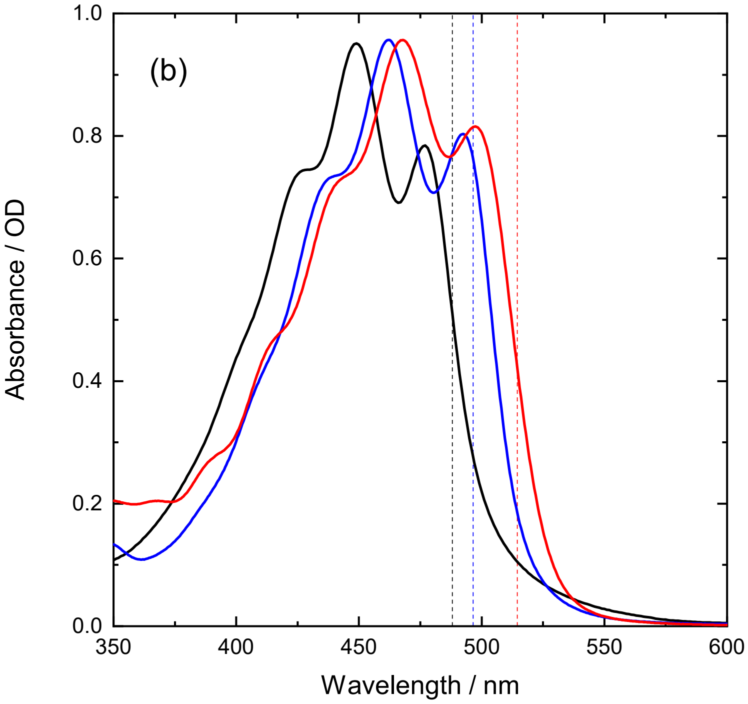


Figure S1: Chemical structures (a) and absorption spectra (b) of the FCP carotenoids (black – Fx in *n*-hexane; blue – DD in pyridine; red – DT in pyridine). Dotted lines indicate the wavelengths of excitation lines used to excite the Raman spectra in Fig. 1.

**Detailed analysis of the resonance Raman changes observed at 496.5 nm excitation occurring during LL-FCPa aggregation**

As stated in the main text, the changes observed in the resonance Raman spectra of LL-FCPa during its successive aggregation are particularly complex to describe in full. This is demonstrated in Figure S2 for 496.5 nm excitation, where the successive differences were computed between the spectrum at each aggregation step (dialysis pH 7.4 & pH 5, and in the presence of Biobeads) and that of trimeric LL-FCPa.

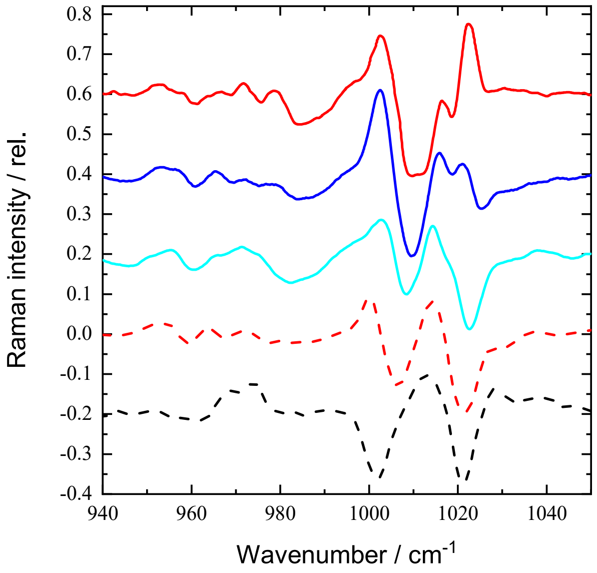


Figure S2: Calculated RR difference spectra for aggregated-minus-disaggregated (trimeric) LL-FCPa at 496.5 nm excitation (see Fig. 2), after normalization on the $\nu_3$ peak. Successive degrees of aggregation were achieved by dialysis at pH 7.4 & 5.5, and by biobead treatment (cyan & blue, and red, respectively). Also shown in dashed lines are the equivalent difference spectra for pH 7.4 dialysis of HL-FCPa, at 501.7 & 514.5 nm excitation (red, black respectively).

It is difficult to disentangle the complexity of this spectral evolution on the basis of these simple difference spectra, which are highly sensitive to the spectral alignment on the wavenumber (X-) axis. Indeed, a small relative shift of the spectra along this axis induces large changes in the shape of the obtained differences, in particular where large intensity bands contribute. We have applied two-dimensional correlation spectroscopy (2DCOS) to this series of spectra, in order to optimise their analysis. 2DCOS is a method designed to analyse the evolution of spectral signals produced by a perturbation[38]. Briefly, it consists in calculating the cross-correlation between a series of spectra according to a variable (in this case, the aggregation state), and yields a quantitative measure of the similarity and dissimilarity of the observed spectral intensity variations. It results in two-dimensional contour maps, with axes expressed in the units of the analysed spectra[38]. One of these maps is termed the synchronous 2D correlation spectrum, and yields, at each point, a measure of the correlation between different parts of the spectrum, *i.e.*, to what degree the intensity at each wavenumber changes simultaneously with the intensity at another wavenumber. Peaks along the diagonal (the autocorrelation spectrum) reveal where the spectrum changes, while corresponding positive and negative cross-peaks aligned with them reflect correlated and anticorrelated changes, respectively (whether their intensity changes in the same direction).

The second map is called the asynchronous 2D correlation spectrum, and describes how much the changes at two spectral points are out of phase – *i.e.,* the degree to which changes at one point are delayed in time with respect to those at another point, according to the evolution of the studied perturbation. In such an asynchronous map, for correlated bands, a positive peak indicates a positive phase shift – the change at the first spectral point (X-axis value) occurs before the change at the second point (Y-axis value). Conversely, a negative peak indicates that the change at the first point occurs after that at the second point. For anticorrelated bands, on the other hand, the signs are reversed – a positive peak indicates a negative phase shift, and *vice versa*. The synchronous map is intrinsically symmetrical, while the asynchronous one is antisymmetrical, relative to the diagonal of the plot. The use of 2D correlation has been spreading in vibrational spectroscopy, and has found many applications[39,40].

Figure S3 shows the 2DCOS synchronous and asynchronous maps, calculated using the Matlab toolbox mat2dcorr[41], for the four spectra obtained at 496.5 nm during successive LL-FCPa aggregation (Fig. 2a). As differences, such maps may be sensitive to the spectral precision along the wavenumber axis. However, the conclusions presented below are not significantly affected by small shifts in the Raman spectra (not shown).

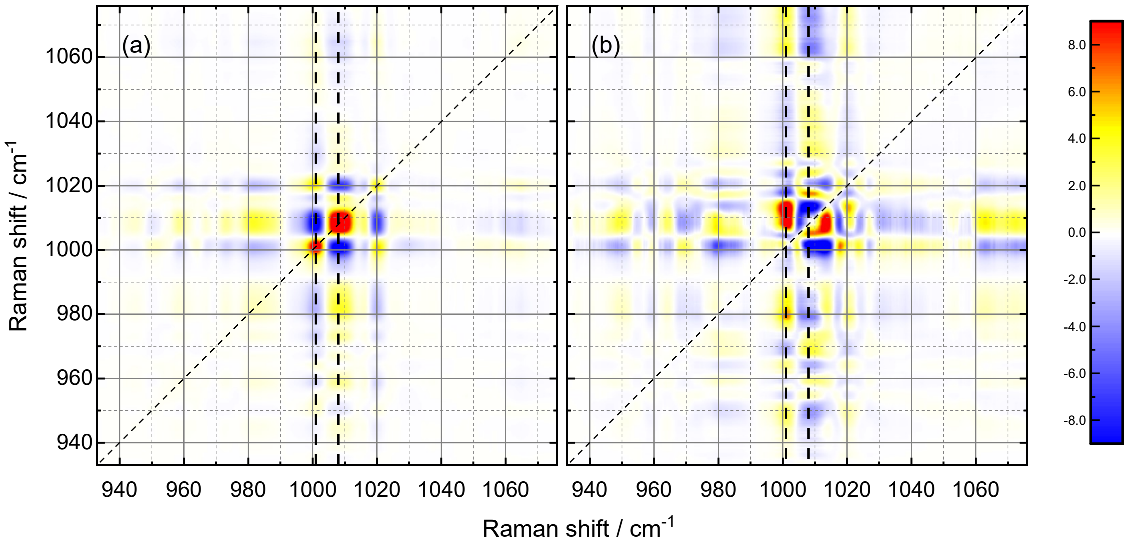


Figure S3: Synchronous (a) and asynchronous (b) 2D maps obtained after cross-correlation of the four spectra displayed in Figure 2a (main text). The average of the four original spectra was used as a reference spectrum. The scale bar represents $\times 10^{-3}$ & $\times 10^{-4}$ relative intensity for (a) & (b), respectively.

The positive cross-peak at position 1001/1020 $cm^{-1}$ in Fig S3a is an example of two bands changing in a correlated way, while the negative cross-peak at 1008/1020 $cm^{-1}$ indicates that these latter two bands change in an anticorrelated way (see dotted lines at X = 1001 & 1008 $cm^{-1}$). The line through coordinate 1001 $cm^{-1}$ coincides successively with a negative peak at ~ 980 $cm^{-1}$, (a positive one with itself at 1001 $cm^{-1}$), a negative one at ~1008 $cm^{-1}$ , and a positive one at ~1020 $cm^{-1}$. Thus the bands at ~980 & ~1008 $cm^{-1}$ are correlated, as are those at ~1001 & ~1020 $cm^{-1}$, whereas the 980/1008 pair is anticorrelated with 1001/1020. Such types of correlations occur between every two components of the spectra, indicating that each perturbation affects, with different amplitudes, all the bands present in the $\nu_3/\nu_4$ regions, and contributions can be formally determined which would otherwise be difficult to observe – such as that at 1008 $cm^{-1}$ in the previous example.

The asynchronous map is even richer. Firstly, it shows that some bands display a complex behaviour, with components changing at different moments during aggregation. This is in particular the case for the bands at 980 $cm^{-1}$, between 1006 and 1018 $cm^{-1}$, and at *ca* 1020 $cm^{-1}$. The shape of the asynchronous cross-peaks along a line at 1001 $cm^{-1}$ is highly distorted as compared to the actual shape of the Raman bands, revealing the presence of various independently-varying components within these bands. Secondly, the sign of the cross-peaks in the synchronous and asynchronous maps determines the order in which spectral changes in different spectral regions occur, *i.e.*, it determines the minimum number of events necessary to fully account for the ensemble of observed spectral variations. Let us determine the order of events involving changes at 980, 1001, 1008, and 1020 $cm^{-1}$. Consider the vertical dashed line through 1008 $cm^{-1}$ on the X-axis - along this line, we observe negative cross-peaks at the other three wavenumbers, viz., 980, 1001, and 1020 $cm^{-1}$. (There are additional positive cross-peaks at 1005 and 1018 $cm^{-1}$ and a strong negative cross-peak at 1013 $cm^{-1}$. The spectral changes at 1018 $cm^{-1}$ may be considered an additional event, but the 1008/1005 and 1008/1018 cross-peaks have negligible intensities in the synchronous map, corresponding to the negligible changes occurring at 1013 & 1018 $cm^{-1}$ in Fig. 2a.) In the synchronous map, 1008 $cm^{-1}$ is correlated with 980 $cm^{-1}$ and anticorrelated with 1001 and 1020 $cm^{-1}$. This indicates that the changes at 1008 $cm^{-1}$ occur after the changes at 980 $cm^{-1}$ and before the changes at 1001 and 1020 $cm^{-1}$. Similarly, the 1001/1020 cross-peak is positive in the synchronous map but negative in the asynchronous map, indicating that the changes at 1001 $cm^{-1}$ are the slowest. The order of events is therefore first at 980 $cm^{-1}$, then at 1008 $cm^{-1}$, followed by 1020 $cm^{-1}$, and lastly at 1001 $cm^{-1}$. Thus a minimum of four spectral events, differentiated temporally, are necessary to fully explain these maps - *i.e.* at least five spectral signatures, including the initial (trimeric) state.
